# Clathrins are involved in the endocytosis of host cytosol in the malaria parasite

**DOI:** 10.1101/2024.12.17.629030

**Authors:** Jun Miao, Gang Ning, Xian Xia, Xiaoying Liang, Amuza Byaruhanga Lucky, Faiza Siddiqui, Hui Min, Chengqi Wang, Xiaolian Li, Z. Hong Zhou, Liwang Cui

## Abstract

In eukaryotic cells, clathrins interact with adaptor protein complexes to regulate key intracellular trafficking events. Specifically, they are associated with the adaptor protein (AP) complex-2 (AP2) to facilitate endocytosis and AP1 to mediate secretion and trafficking between endosome and Golgi. In *Plasmodium falciparum,* recent studies revealed that the Kelch domain-containing protein 13 and AP2 participate in hemoglobin uptake via cytostomes. However, clathrins appear not to be involved in this process because they primarily associate with AP1. To investigate the roles of clathrins in *P. falciparum*, we characterized the clathrin heavy chain (*Pf*CHC), the clathrin light chain (*Pf*CLC), and the AP1 γ subunit (*Pf*AP1 γ). Extensive interactome analyses confirmed the major association of clathrins with AP1 components alongside proteins involved in cytostome formation. Live-cell imaging and protein colocalization studies showed that *Pf*CHC, *Pf*CLC, and *Pf*AP1 γ are localized in the parasite cytoplasm, predominantly at the parasite periphery and near the cis-Golgi. Ultrastructural studies using ascorbate peroxidase 2- based electron microscopy confirmed their presence at coated vesicle-like structures at the parasite periphery and, unexpectedly, at the collars of cytostomes. Consistent with these observations, the knockdown of *Pf*CHC led to the formation of abnormally long cytostome tubes and impaired hemoglobin digestion. This study demonstrates that clathrins are essential for proper cytostome formation in *P. falciparum*, highlighting their critical role in the parasite’s hemoglobin uptake and digestion processes.

**IMPORTANCE:** Malaria is still one of the most severe public health problems worldwide and understanding how malaria parasites obtain hemoglobin from their host cells, red blood cells (RBCs), is critical for identifying targets for antimalarials. In this study, affinity purification revealed that clathrins (*Pf*CHC and *Pf*CLC) were not only mainly associated with AP1 complex but also marginally interacted with proteins that participated in hemoglobin uptake. Localization analysis demonstrated that clathrins and AP1 coated the clusters of vesicles at the parasite periphery and at the neck of the cytostome, an organelle for hemoglobin uptake. Consistently, the knockdown of *Pf*CHC caused the formation of abnormally long cytostome tubes and reduced hemoglobin digestion. Collectively, this study demonstrates that clathrins play a critical role in the parasite’s hemoglobin uptake.

## INTRODUCTION

Clathrin-coated vesicles (CCVs) in eukaryotes are well-established (1–6). Triskelia, composed of three clathrin heavy chains (CHCs) and three clathrin light chains (CLCs), are the assembly units of the polygonal lattice that make up clathrin-coated vesicles (2, 7). The most abundant proteins in the clathrin coat are the heterotetrameric adaptor protein (AP) complexes. Each AP complex contains two large subunits (α/γ/δ/ε/ζ and β1-5) of ∼100 kDa, a medium subunit (μ1-5) of ∼50 kDa, and a small subunit (σ1-5) of ∼20 kDa. APs are classified into five types (AP1-5). AP1 involved traffic from the TGN to the endosome or polarized plasma membrane for secretion and the AP2 located at the plasma membrane for endocytosis (2, 7–11). While AP3 also plays a role in the traffic between the Golgi and endosomes, AP4 and AP5 are not associated with coated vesicles (2, 12). Different sorting signals present in the cargo molecules are recognized by different subunits of AP complexes (2, 12).

Of the five malaria species infecting humans, *Plasmodium falciparum* causes the most severe malaria and is responsible for nearly half a million deaths annually (13). The *P. falciparum* genome encodes most components of clathrin complexes (14, 15). Recent studies found that *Pf*CHC, *Pf*AP1 μ1, and *Pf*AP2 μ2 were localized in the cytosol of the parasite, and *Pf*AP2 μ2 is essential for schizont maturation (16–18). Yet, the significance of CCVs in the malaria parasites has not been investigated.

Apicomplexan parasites take up the host cytoplasm by endocytosis via a structure called micropore, which comprises parasite plasma membrane (PPM) invagination surrounded by electron-dense rings at its collar (19–21). Type I micropore occurs in openings of the inner membrane complex (IMC) of *Plasmodium* sporozoite and *T. gondii*. In *Plasmodium,* the cytostome (also called type II micropore) has an additional parasitophorous vacuolar membrane invagination but without the IMC restriction and is specific for the uptake of host cell hemoglobin (Hb) (19). In *P. falciparum*, Kelch domain-containing protein 13 (PfK13), the key determinant of artemisinin resistance (17, 18, 21–24), and *Pf*AP2 complex-associated proteins were localized at the cytostome collar (17, 18, 22, 24, 25). Similar localization patterns of K13 and AP2 subunits were confirmed in *T. gondii* (26, 27). These findings suggest the involvement of K13 and AP2 complex in endocytosis in both *P. falciparum* and *T. gondii*. Consistently, knockdown (KD) of K13 and its associated proteins in both species disturbed the formation of the cytostome or micropore and affected the uptake of Hb or host cell materials (18, 24, 26, 27). Unexpectedly, clathrins were not associated with the *Pf*AP2 complex, suggesting that they do not play a major role in endocytosis. Instead, *Pf*CHC was mainly associated with the AP1 complex in *P. falciparum* (17, 18). Therefore, clathrins may play parasite-specific roles in the malaria parasite.

In this study, the unique features of clathrins and their predominant association with the AP1 complex in *P. falciparum* prompted us to characterize their functions in vesicle transport. Through extensive interactome, localization, and gene manipulation studies, we surprisingly found that clathrins and AP1 were involved in endocytosis of Hb via mediating cytostome processes.

## RESULTS

### *P. falciparum* encodes conserved clathrins and AP1 complex

While the *Pf*CHC (PF3D7_1219100) has a conserved domain structure organization with ∼40% identity to CHCs in model eukaryotic organisms, it contains five Asn-rich insertions longer than 28 aa, accounting for an overall increase of protein size by >300 amino acids (aa) (**Fig. S1A, S1B**). Despite these insertions, the region predicted to interact with the CLC and residues (Q89, F91, K96, and K98) critical for β-arrestin and β-adaptin binding in the N-terminal β-propeller domain in vertebrate CHCs (28–31) are perfectly conserved in *Pf*CHC. *Pf*CLC (PF3D7_1435500) is also conserved in domain organization but only has ∼20% identity to vertebrate CLCs. In the consensus region, the acidic patch (DEG vs. EED in human CLC) and residues (W153 and I175 in *Pf*CLC vs. W105 and W127 in human CLC) critical for binding CHC (32, 33) are relatively conserved (**Fig. S1C, S1D**). *Pf*AP1 γ subunit (PF3D7_1455500) has ∼30% identity to vertebrate AP1 γ1s with a 99-aa insertion in the trunk domain and a longer unstructured linker (**Fig. S1E, S1F**). The other AP1 subunits, *Pf*AP1 β1, μ1, and σ1, display higher identities (∼48%, ∼60%, and ∼57%) to their respective vertebrate counterparts, respectively (**Fig. S1G-L**). *Pf*CHC, *Pf*CLC, and *Pf*AP1 subunits show the highest levels of sequence identity to their respective counterparts from *T. gondii* (**Fig. S1**). These results indicate that *P. falciparum* clathrins harbor both conserved and diverse domain structures.

### Affinity pulldowns identify proteins involved in cytostome formation

Previous reports showed that *Pf*CHC was mainly associated with AP1 subunits by proximity labeling (BioID) and affinity purification (17, 18). To confirm these findings, we tagged the endogenous *Pf*CHC, *Pf*CLC, and *Pf*AP1 γ with GFP using a single-crossover recombination strategy (**Fig. S2A, S2B**). Western blots with anti-GFP antibodies revealed ∼260, ∼60, and ∼150 kDa bands, consistent with the molecular weights predicted from the GFP-fused *Pf*CHC, *Pf*CLC, and *Pf*AP1 γ, respectively (**Fig. S2C-E**). The expression levels of these proteins were increased during the intraerythrocytic developmental cycle (IDC) (**Fig. S2C-E**).

We first precipitated proteins from trophozoites of the *Pf*CHC::GFP parasite line using GFP-trap agarose. Using liquid chromatography and tandem mass spectrometry (LC-MS/MS), we identified 87 proteins using stringent parameters by Significance Analysis of INTeractome (SAINT) analysis (a threshold of probability above 90% and false discovery rate (FDR) below 2.5%) (34) (**Table S1**). *Pf*CHC, *Pf*CLC, and AP1 subunits were identified at high abundance, confirming that *Pf*CHC is principally associated with AP1 (**Table S1**). Comparing this interactome result with the 40 and 63 *Pf*CHC-associated proteins identified previously by BioID and anti-GFP magnetic beads, respectively (17, 18), we identified nine shared proteins including *Pf*CHC, *Pf*CLC, AP1 subunits, Sortilin (an escort protein for trafficking) (35, 36), a putative AP4 complex accessory subunit Tepsin (**Table S1**). Like the previous assumption (18), the identification of Tepsin indicates that clathrins may be associated with the AP4 complex.

To further validate these results, we conducted reciprocal GFP-trap pulldowns from the *Pf*CLC::GFP parasites. In three replicates, we identified 332 proteins using the same cutoffs for the SAINT analysis (**Table S2**). Consistently, *Pf*CHC, *Pf*CLC, AP1, Sortilin, and Tepsin were identified at high abundance. Additionally, four AP4 subunits, two AP2 subunits (AP2 μ2 and AP2 α), and two AP3 subunits (AP3 β and AP3 μ3) were also identified at lower abundance (**Fig. 1**, **Table S2**). Furthermore, some proteins related to Hb digestion, including three aminopeptidases and falstatin, a cysteine protease inhibitor, were also identified from the *Pf*CLC pulldowns (**Fig. 1**, **Table S2**). Interestingly, we also identified five subunits of the V-type proton ATPase, which was localized at the food vacuole (FV) and PPM and is involved in Hb uptake (37, 38) (**Fig. 1**, **Table S2**). Taken together, these interactome analyses revealed that clathrins were also associated with other AP (AP2-4) and proteins involved in Hb uptake and digestion.

**Figure 1.**
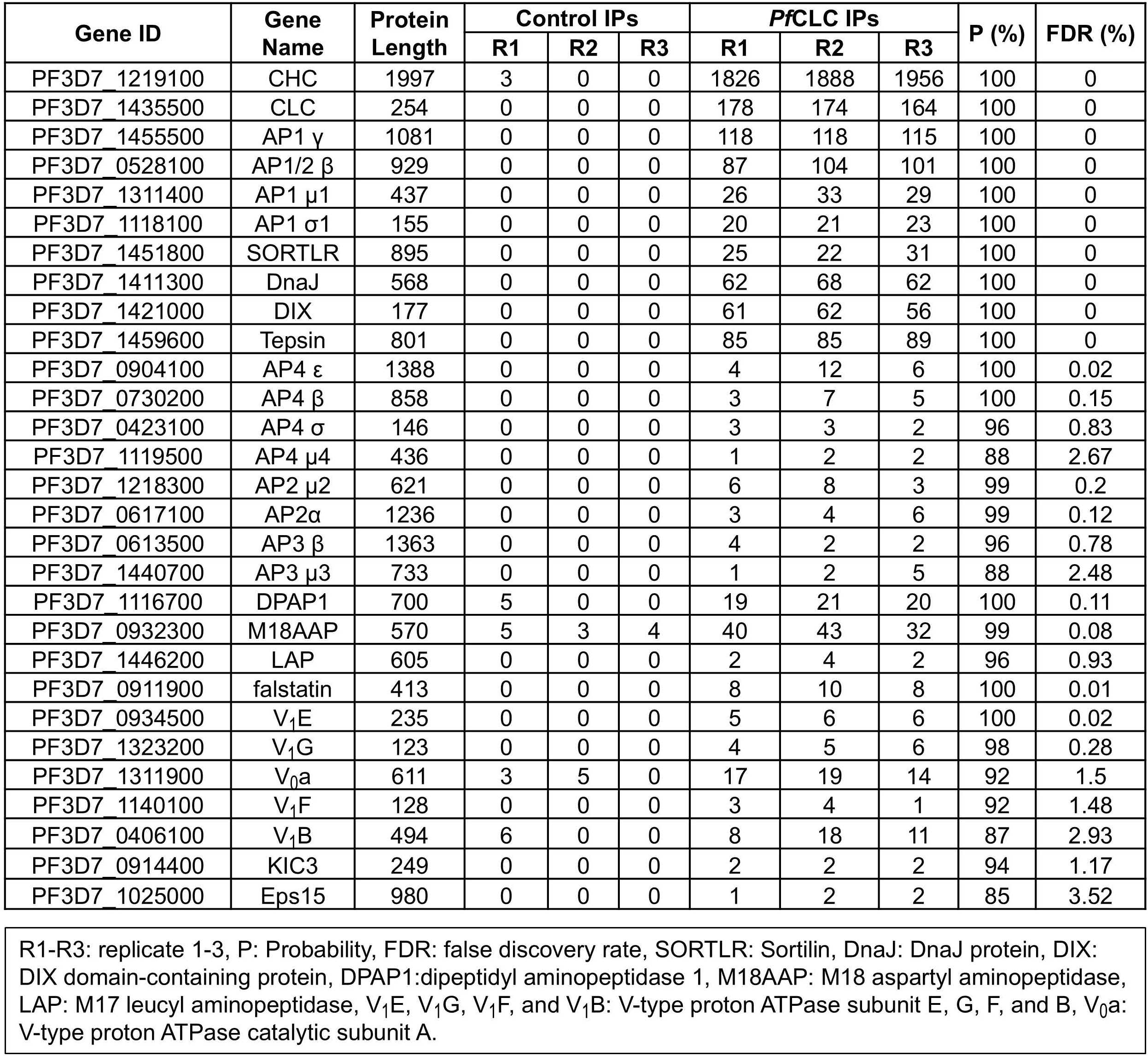
List of the important proteins identified from *Pf*CLC::GFP parasites by GFP- trap. R1-R3: replicate 1-3, P: Probability, FDR: false discovery rate, SORTLR: Sortilin, DnaJ: DnaJ protein, DIX: DIX domain-containing protein, DPAP1:dipeptidyl aminopeptidase 1, M18AAP: M18 aspartyl aminopeptidase, LAP: M17 leucyl aminopeptidase, V1E, V1G, V1F, and V1B: V- type proton ATPase subunit E, G, F, and B, V0a: V-type proton ATPase catalytic subunit A.

To confirm the interaction between clathrins and AP1, we conducted reciprocal GFP-trap pulldowns from the *Pf*AP1 γ::GFP parasite and identified 342 interacting proteins (**Table S3**). As expected, *Pf*CHC, *Pf*CLC, and AP1 were highly abundant in the pulldowns (**Table S3**).

Furthermore, two aminopeptidases (M18 aspartyl aminopeptidase and M1-family alanyl aminopeptidase) and the five subunits of the V-type proton ATPase were also detected (**Table S3**). Interestingly, the interactomes of *Pf*CLC and *Pf*AP1 γ also shared KIC3 and Eps15 with the *Pf*K13 interactome identified by BioID (18) although these proteins were relatively scarce in *Pf*CLC and *Pf*AP1 γ pulldowns (**Fig. 1**, **Table S2, S3**), suggesting that the *Pf*K13 complex may be loosely or indirectly associated with the clathrins-AP1.

### Clathrins are localized at the parasite periphery and the collar of the cytostome

To understand the potential involvement of clathrins in vesicle transport in *P. falciparum*, we first evaluated their localizations in the three GFP-labeled parasite lines. Live-cell fluorescence microscopy revealed the presence of large fluorescent puncta in the parasite, with the size and number of GFP puncta substantially increasing during the IDC (**Fig. 2A-C**). Under structured illumination microscopy (SIM), the GFP puncta in the parasite appeared as spheres, teardrops, or discs with variable sizes of 100–300 nm in diameter (**Fig. 2D**, **S3**). Intriguingly, these GFP puncta were localized at the periphery of trophozoites (**Fig. 2D**, **S3**). Indirect immunofluorescence assay (IFA) revealed that *Pf*CHC-GFP was juxtaposed with the cis-Golgi marker ERD2 (**Fig. 2E**, ∼23% colocalization) and was within the PV marker Exp2 that defines the parasite boundary (**Fig. 2F**).

**Figure 2.**
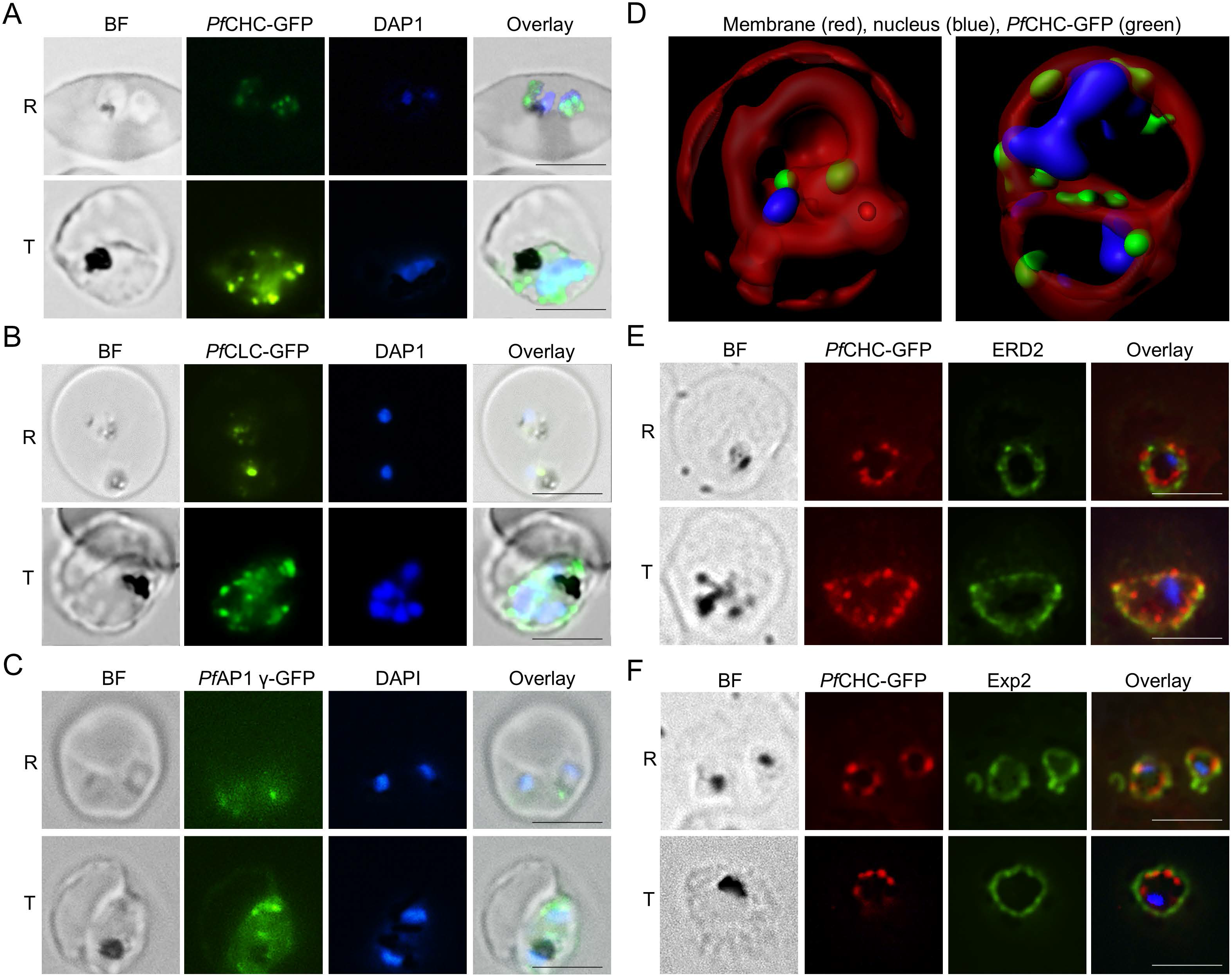
The localization of clathrins and AP1 γ in *P. falciparum* by IFA. **A-C**. Live imaging shows GFP foci in the *Pf*CHC::GFP (**A**), *Pf*CLC::GFP (**B**), and *Pf*AP1 γ::GFP (**C**) parasite lines during the asexual development. BF: bright field, R: ring, T: trophozoite. The size of the scale bar is 5 μm. **D**. Two representative images of *Pf*CHC::GFP infected erythrocytes by structure illumination microscopy showing the three-dimensional display of cells with Bodipy membrane staining (GFP foci in green, nucleus in blue, and membrane in red). **E** and **F**. Representative images show the partial colocalization of *Pf*CHC with Golgi marker ERD2 (**E**) and PV marker Exp2 (**F**) in the asexual stage parasites. The size of the scale bar is 5 μm.

To refine the ultrastructures of peripherally localized clathrin puncta, we used the ascorbate peroxidase 2 (APEX2)-based transmission electron microscopy (TEM). APEX2 is an engineered ascorbate peroxidase that can convert diaminobenzidine (DAB) into an insoluble polymer in the presence of H2O2 for a short time, which shows strong EM contrast at the site of the APEX2-tagged protein after staining with osmium tetroxide (OsO4) (39). The central advantage of this technology is that APEX2 remains active after glutaraldehyde fixation to offer excellent preservation of subcellular structures. The DAB polymer remains tightly localized to the site of production and does not cross membranes, thus allowing precise localization of the labeled protein (39, 40). Using the single crossover homologous recombination strategy, we tagged the C-termini of the endogenous *Pf*CHC, *Pf*CLC, and *Pf*AP1 γ with APEX2 (**Fig. S4**).

APEX2-based TEM analysis in the *Pf*CHC::APEX2 parasite line revealed that the electron-dense materials coated the clusters of the vesicle-like structures in the parasites and near PPM (**Fig. 2A**, white arrows). Surprisingly, electron-dense materials also coated the cytostome collars (**Fig. 3B, 3C**, black arrows). While most of the cytostomes were vertically dissected in the sections, showing two layers of the electron-dense materials at the cytostome collars (**Fig. 3B**), some were horizontally cut at the cytostome neck, displaying electron-dense material- labeled double-rings (**Fig. 3C**, black arrows). Furthermore, the electron-dense coats of the vesicle-like structures were often localized near the cytostomes (**Fig. 3C**, white arrows). In contrast, the wild-type parasite 3D7 under the same APEX2 reaction (DAB and H2O2) and OsO4 staining did not show any electron-dense materials at the parasite periphery and cytostome collar (**Fig. 3D**, black arrows). The same patterns of electron-dense materials were also identified in the *Pf*CLC::APEX2 parasite line (**Fig. S5**).

**Figure 3.**
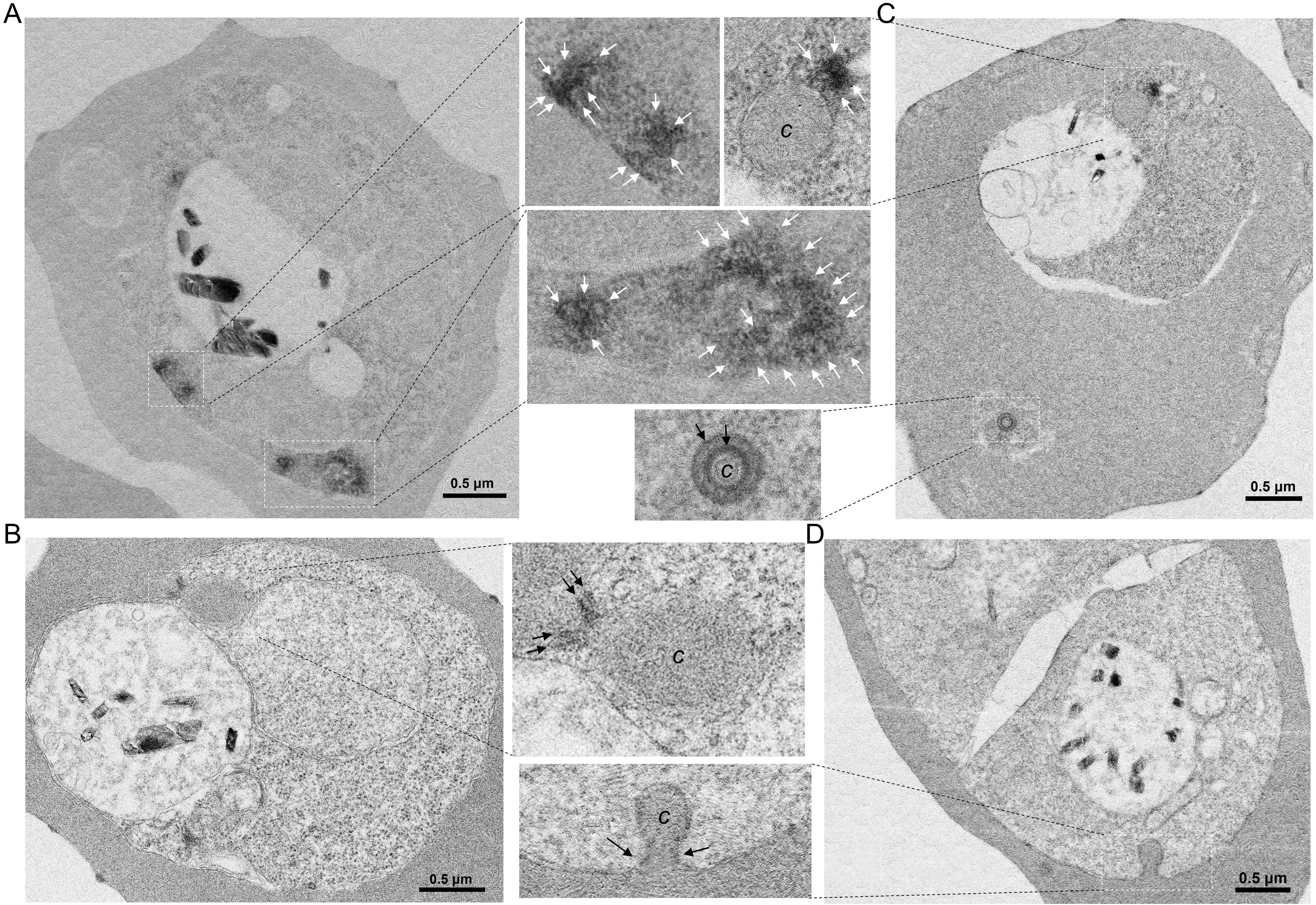
The localization of *Pf*CHC by APEX2-based EM. **A**. A representative EM image shows that *Pf*CHC is localized in vesicle-like structures in the parasite cytosol (white arrows). **B**. A representative image displays that *Pf*CHC is localized at the cytostome neck (black arrows). *C*: cytostome. **C**. A representative EM image indicates that *Pf*CHC is localized in vesicle-like structures in the parasite cytosol (white arrows) close to the cytostome and at the collar of a horizontally dissected cytostome (black arrows). **D**. A representative EM image shows electron-dense materials were detected neither at a cytostome neck (black arrows) nor the parasite periphery of the non-APEX2 tagged parasites (3D7) with the same APEX2-based EM conditions (OsO4statining after reaction (DAB+H2O2).

We then performed APEX2-based TEM analysis in the *Pf*AP1 γ::APEX2 parasite line. Like the localization of *Pf*CHC and *Pf*CLC, *Pf*AP1 γ was also localized at vesicle-like structures near PPM (**Fig. 4A**, **3B**, **S6A-S6C**, white arrows). In contrast to the double-layers of *Pf*CHC and *Pf*CLC coats at the cytostome necks, only a single layer of electron-dense materials was detected at the cytostome collars in the *Pf*AP1 γ::APEX2 parasite line when the cytostomes were dissected vertically (**Fig. 4A, 4B, 4D, 4E**, **S6 E,** black arrows) or horizontally (**Fig. 4C S6C, S6D**, black arrows). Likewise, the electron-dense materials-coated vesicles were often localized near the cytostomes (**Fig. 4D, 4E, S6E**, white arrows). Additionally, the electron-dense materials were found to coat the entire surface of small invagination at PPM (**Fig. 4F, 4G**). Two parallel, electron-dense material-coated invaginations in Fig. 3F were reminiscent of the clusters of cytostomes observed by ultrastructural expansion microscopy (25), indicating that these are likely to be early-stage cytostomes.

**Figure 4.**
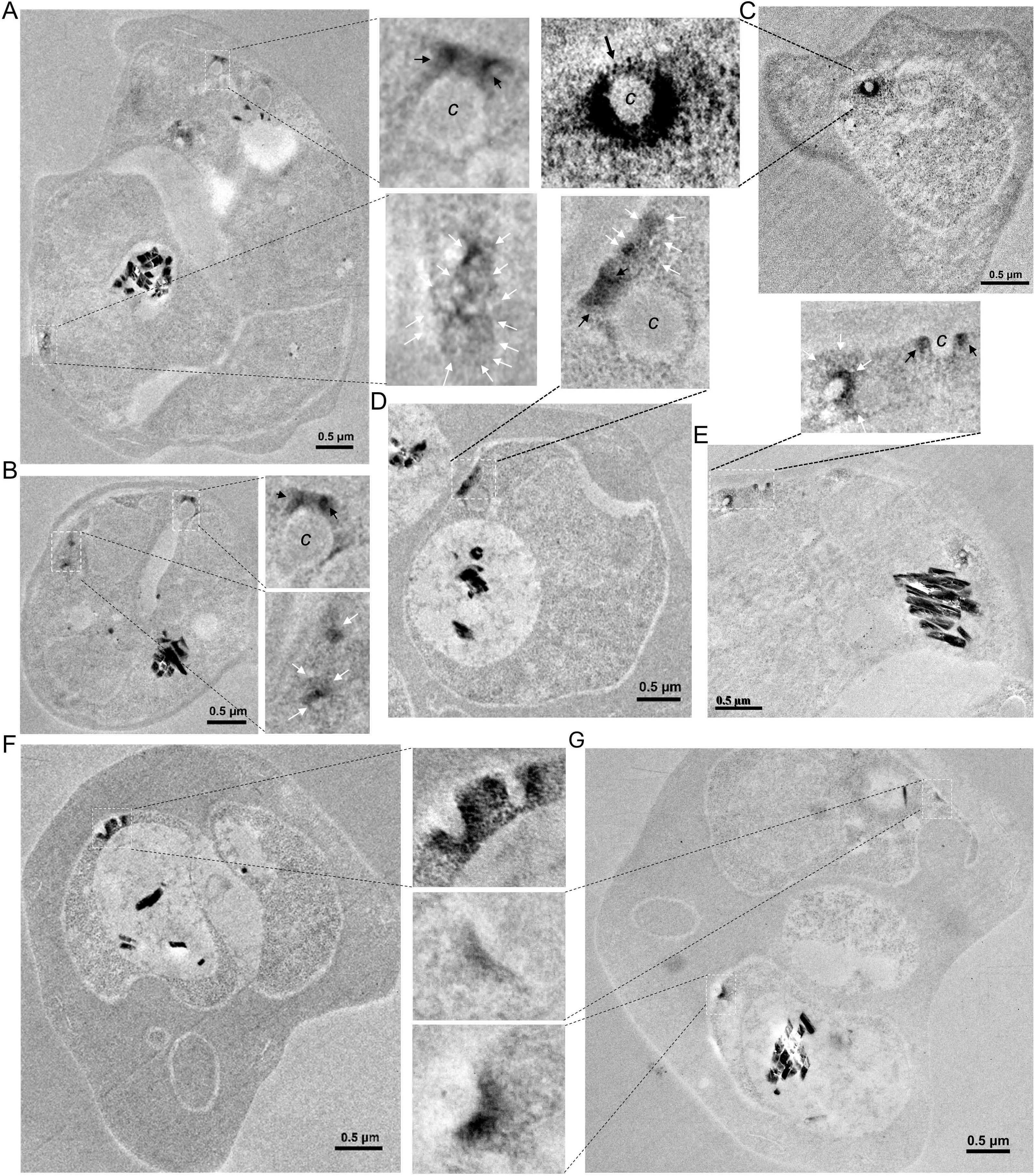
The localization of *Pf*AP1 γ by APEX2-based EM. **A** and **B**. Two representative EM images show that *Pf*AP1 γ is localized in vesicle-like structures in the parasite cytosol (white arrows) and at the cytostome neck (black arrows). *C*: cytostome. **C**. A representative image indicates a horizontally dissected cytostome with a single layer of electron-dense materials at its collar. **D** and **E**. Two representative EM images display that *Pf*AP1 γ is localized in vesicle-like structures in the parasite cytosol close to cytostomes (white arrows) and at the collar of cytostome (black arrows). **F** and **G**. Two EM images show that the electron-dense materials coat the outer surface of two parallel small invaginations (early-stage cytostomes) (**F**) and two separated small invaginations (**G**) from PPM.

### CCVs in malaria parasites appear more diverse

APEX2-based EM easily identified electron-dense material-coated vesicle-like structures in the parasites (**Fig. 3, 4, S5, S6**). This is in stark contrast to the rare report of CCVs in the parasites (41). To further elucidate the structure of CCVs in *P. falciparum*, we used two approaches. First, we purified CCVs from the *Pf*CHC::GFP parasite using anti-GFP magnetic beads followed by negative staining TEM analysis. We detected vesicles with conspicuous coat structures, many similar in size to canonical CCVs, while others were much larger and pleomorphic (**Fig. 5A**, **S7A)**. The thickness of the electron-dense coat is ∼7 nm, significantly thinner than that in CCVs of mammalian cells (∼15 nm) (7, 42). This coat could be mistaken for double membrane-bound vesicles (**Fig. S7B**). Secondly, we harvested CCVs from wild-type parasites by sucrose/Ficoll-gradient ultracentrifugation, a technique commonly used to prepare homogeneous CCVs from mammalian cells (43). TEM and cryoEM revealed typical CCV-like vesicles with an average size of 36.9 nm (30–50 nm, n=104) in the harvested materials (**Fig. 5B, 5C**). Analysis of the harvested materials by LC-MS/MS detected many proteins including *Pf*CHC, *Pf*CLC, and *Pf*AP1 subunits at high abundance (**Table S4**).

**Figure 5.** Morphology of CCVs in *P. falciparum*. **A.** TEM image of CCVs purified by GFP-trap beads from *Pf*CHC::GFP parasites. Two inserts show two smaller CCVs. **B** and **C.** TEM (**B**) and Cryo-EM (**C**) images of CCVs purified by ultracentrifugation from wild-type 3D7 parasite after negative staining (**B**) and cryofixation (**C**).

**Figure 6.**
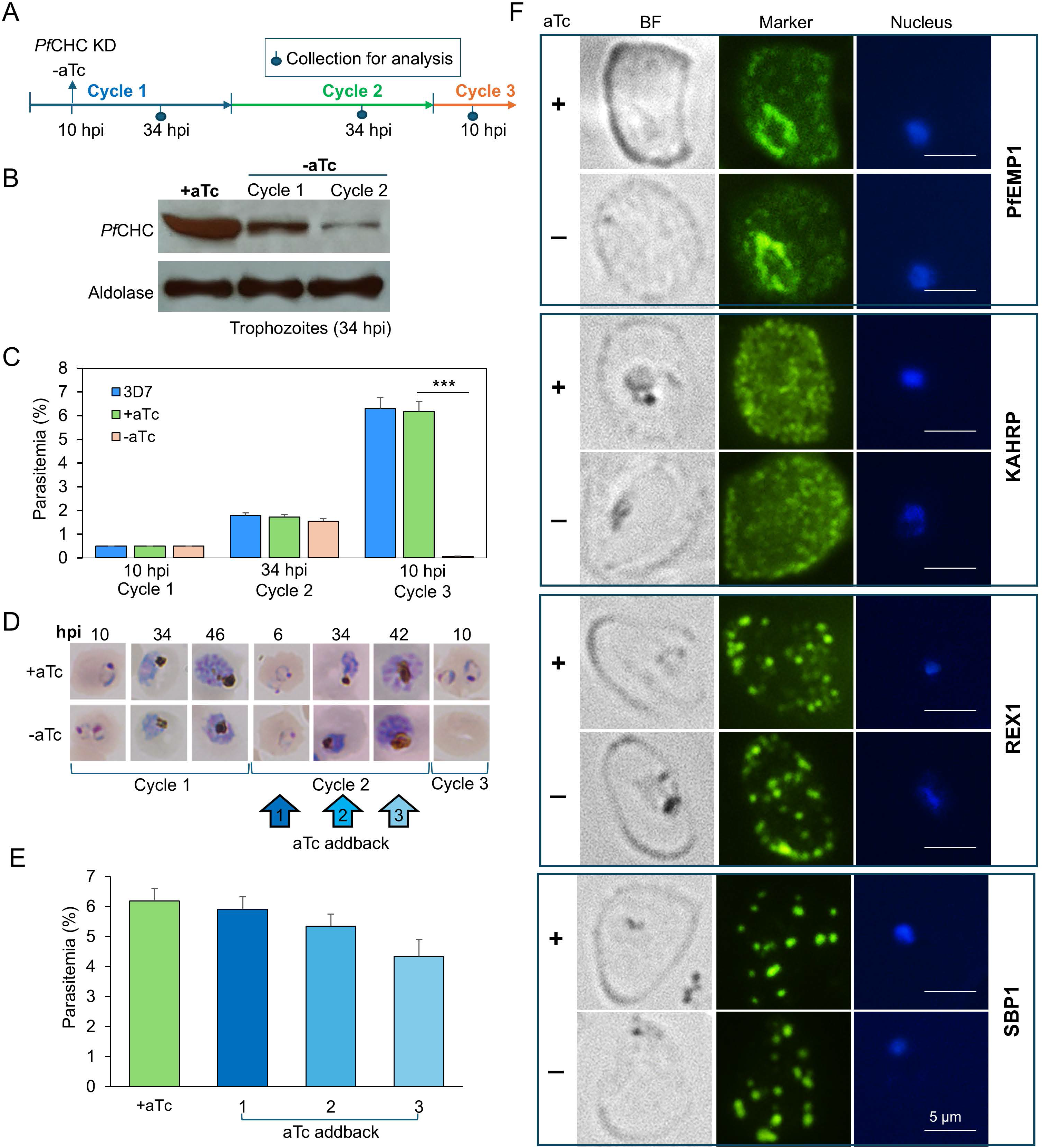
Growth phenotypes and protein export upon *Pf*CHC KD. **A**. A schematic diagram shows the experimental workflow on the analysis of *Pf*CHC expression (B) and growth (**C**-**E**) after *Pf*CHC KD. **B**. Western blot with anti-GFP antibodies showed the progressive reduction of *Pf*CHC-GFP protein level in cycle 1 (∼60%) and cycle 2 (∼90%). aTc was withdrawn from the culture at 10 h post-invasion (hpi). The intensities of protein bands were measured using ImageJ. **C** and **D**. Parasite growth after *Pf*CHC KD by measuring parasitemia (C) and checking parasite development by Giemsa staining (**D**). The starting parasite cultures at 0.5% ring grew normally in cycle 2 but no ring-stage parasites appeared at 10 hpi in cycle 3. *** : *P* <0.001. **E**. When aTc was added back to the culture in cycle 2 at the ring, trophozoite, or schizont stage (6, 34, or 42 hpi), a variable percentage of parasites could survive in the next cycle. **F**. IFA with specific antibodies showed the export of three PNEPs (PfEMP1, REX1, and SBP1), and one PEXEL-containing protein (KAHRP) was not disturbed after *Pf*CHC KD for 48h (-aTc) (aTc was withdrawn at early trophozoite stage for 48h) compared to their controls (+aTc). The parasite nuclei were stained by DAPI. BF: bright field. The size of the scale bar is 5 μm.

### *Pf*CHC KD leads to reduced Hb digestion

To understand clathrin function, we attempted to disrupt *PfCHC* by single and double crossover recombination but without success, consistent with the essentiality of this gene from transposon mutagenesis analysis (44). We then applied the TetR-DOZI conditional KD system to reduce the expression of *Pf*CHC-GFP (45, 46) (**Fig. S8A, S8B**). In the engineered parasite TetR- *Pf*CHC::GFP, withdrawing anhydrotetracycline (aTc) from the 10 hour-post invasion (hpi) ring-stage parasites for about 24 h (defined as 34 hpi trophozoites in Cycle 1), *Pf*CHC expression was reduced to ∼40% (**Fig. 6A, 6B**). Culturing the TetR-*Pf*CHC::GFP parasites without aTc for an additional 48 h to the 34 hpi trophozoite stage of Cycle 2 resulted in approximately 90% reduction of *Pf*CHC expression (**Fig. 6A, 6B**). Compared to the TetR-*Pf*CHC::GFP parasite cultured in parallel with constant aTc, parasite growth was only slightly reduced after aTc withdrawal in Cycle 1, and the majority of parasites developed into mature schizonts. However, in the ring stage (10 hpi) of Cycle 3 without aTc, when all schizonts had egressed, we observed a very low number of ring-stage parasites in the culture. In contrast, parasites cultured in parallel with aTc all developed into the ring stage (**Fig. 6C, 6D**). These results suggested that merozoites released from the schizonts of Cycle 2 were defective in the invasion of RBCs. Adding aTc back into the culture at the ring, trophozoite, and schizont stage in Cycle 2 enabled parasite growth into Cycle 3 at ∼95%, ∼86%, and ∼70% efficiency compared with cultures with aTc, confirming that the observed invasion defect was due to *Pf*CHC KD (**Fig. 6D, 6E**). Intriguingly, the later aTc was added back to the culture, the higher levels of gametocytes were generated in the culture (**Fig. S8C**), indicating that *Pf*CHC KD increased commitment to gametocytogenesis.

The location of clathrin-coated vesicle-like structures at the parasite periphery (**Fig. 3, 4,** S5, S6) suggests that clathrins may be involved in protein export into RBCs. To investigate whether *Pf*CHC KD influences this process, we performed IFA on two PEXEL motif-containing proteins (KAHRP and FIKK4.2) and four PEXEL-negative exported proteins (PfEMP1, REX1, REX3, and SBP1) (47, 48). No discernable reduction in the export of these proteins was detected 48 h after aTc withdrawal (**Fig. 6F**, **S8D**).

The localization of clathrins at cytostome collars suggests their potential involvement in endocytosis. In Giemsa-stained smears, we observed that the hemozoin pigment in the food vacuole (FV) in the *Pf*CHC KD parasite was much lighter in color and occupied a smaller area than in the control (**Fig. 7A**, **S8E**), suggesting the accumulation of a lower amount of hemozoin in *Pf*CHC KD parasites. To confirm this observation, we quantified hemozoin from equal numbers of synchronized *Pf*CHC-KD and control parasites at the trophozoite stage. The results showed that *Pf*CHC-KD parasites contained a significantly lower amount of hemozoin than control parasites (**Fig. 7B** *P* < 0.01, t-test). TEM revealed long tube-like cytostomes in the *Pf*CHC KD parasites (-aTc) after withdrawal of aTc for 48 h compared to the Hb-containing small vesicles in the control parasites (+aTc), indicating that *Pf*CHC KD led to a defect in the pinching-off of Hb-containing vesicles from cytostomes and therefore caused less amount of Hb to be delivered to the FV (**Fig. 7C**, **S8F**).

**Figure 7.**
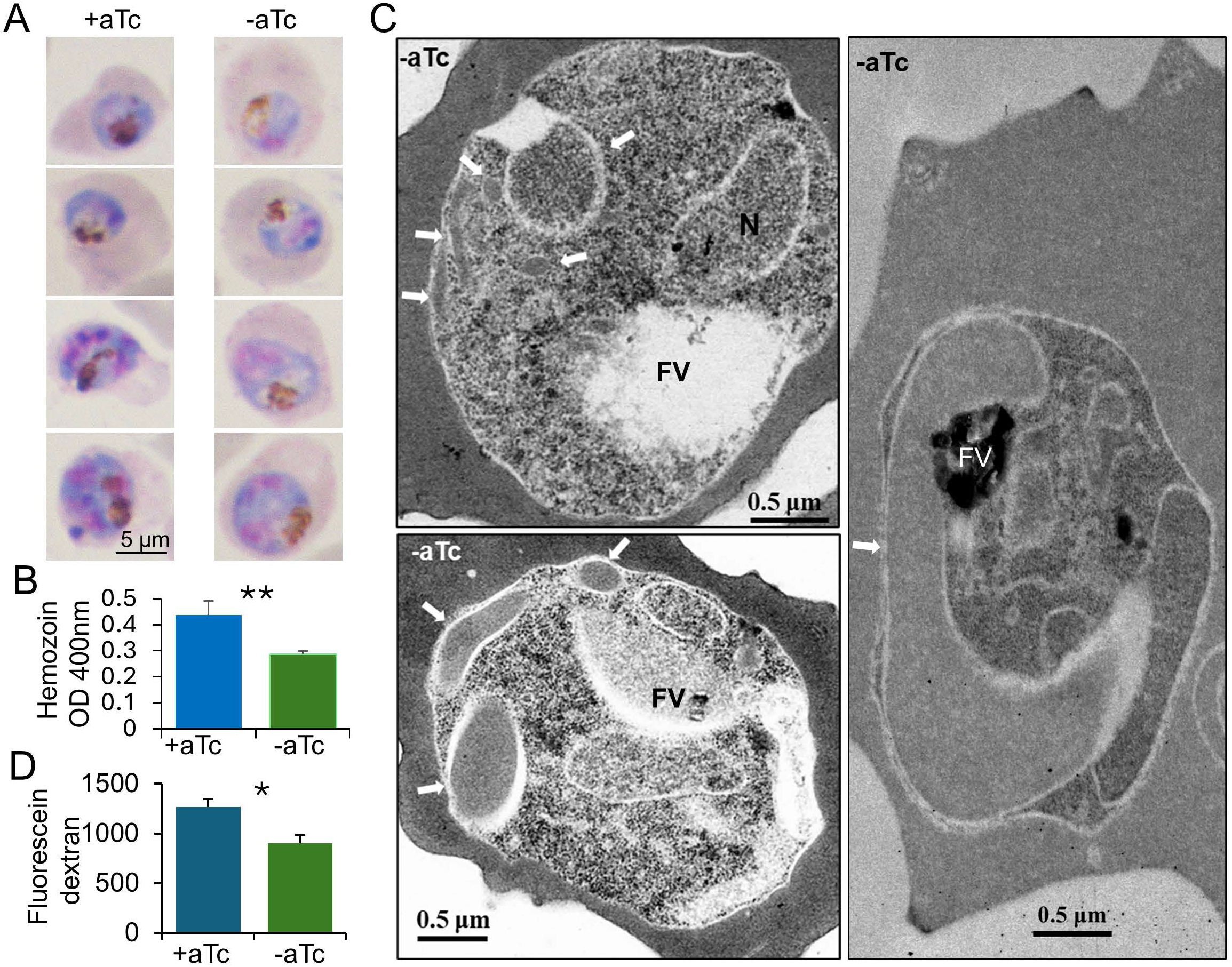
*Pf*CHC KD led to abnormal cytostome and lower Hb digestion. **A**. Representative Giemsa staining images show the hemozoin blocks before (+aTc) and after (- aTc) *Pf*CHC KD. **B**. A bar graph shows hemozoin levels in the synchronized parasites at the trophozoite stage before (+aTc) and after (-aTc) *Pf*CHC KD. Hemozoin was measured by the absorbance at 400 nm. **C**. Three representative EM images show the abnormal cytostome (long tubes) (white arrows) after *Pf*CHC KD (-aTc). FV: food vacuole. **D**. A bar graph displays the overall levels of fluorescein-dextran uptake from resealed RBCs into the trophozoite-stage parasites before (+aTc) and after (-aTc) *Pf*CHC KD. The green fluorescence signals of fluorescein-dextran were measured by flow cytometry.

Since the mouth of the cytostome is open to the RBC cytosol, lysis of the RBC membrane will release Hb from both RBC and cytostomes but not from the Hb-containing vesicles that have been pinched off from cytostomes. Thus, if lysis of RBC membrane releases Hb from abnormally long tube-like cytostomes in the *Pf*CHC KD parasites, the Hb content in the *Pf*CHC KD parasites will be lower than its control. To verify this, resealed RBCs containing fluorescein-dextran were infected by *Pf*CHC KD parasites. Flow cytometry indicated that fluorescence signals in *Pf*CHC KD parasites (-aTc) were significantly lower than their controls (+aTc), suggesting that Hb uptake was reduced upon *Pf*CHC KD (**Fig. 7D** *P* < 0.05, t-test).

## DISCUSSION

In this study, we studied the interactome, localizations, and functions of clathrins in *P. falciparum*. Interactome studies confirmed the association of clathrins mainly with the AP1 complex and, to a much lower extent, with other AP complexes. In addition, we also identified their associations with proteins involved in Hb uptake/digestion. IFA and APEX2-based EM precisely identified the localization of *Pf*CHC, *Pf*CLC, and *Pf*AP1 γ at vesicle-like structures, which were clustered together near the PPM and at the collar of the cytostome. *Pf*CHC KD led to defects in merozoite invasion, the accumulation of abnormal cytostome structures, and consequently reduced Hb digestion.

It is conceivable that *Pf*AP1-based CCVs near the Golgi mediate secretion and trafficking between the Golgi and endosomes in the parasite. Since protein export to the host RBC cytosol was not noticeably altered upon *Pf*CHC KD, *Pf*AP1-containing CCVs may be primarily involved in protein secretion to the PPM or PV for cytostome biogenesis and protein trafficking between the Golgi and endosomes (cytostome and FV). Since Sortilin was also identified in the clathrin IPs from the parasite, clathrin/AP1-based transport may have the conserved function in protein transport among the Golgi, endosome, and PPM (exocytosis) (49–52). Previous reports showed that some Hb proteases, such as falcipain and plasmepsin II, were found to be transported to PV or cytostome *en route* to the FV (53–56). Two recent studies also showed that the V-type ATPase, which regulates the pH of FV by pumping protons and consists of eight cytosolic V1 domain-containing subunits and five membrane-embedded V0 domain-containing subunits, was localized at the FV and PPM (37, 38). KD of its subunits resulted in accumulated Hb-containing vesicles in the FV, indicating that this complex is also potentially transported to the cytostome to create acidic conditions *en route* to the FV (37, 38). Interestingly, previous pulldowns using anti-GFP magnetic beads (17) and our current GFP-trap pulldown studies identified several proteases, a protease inhibitor (falstatin), and subunits of the V-type ATPase, suggesting that they may be cargoes of AP1-based CCVs.

Conventional AP2-based CCVs in eukaryotes mediate endocytosis. Therefore, there has been long-standing speculation that AP2-based CCVs might mediate endocytosis via the cytostome in malaria parasites (17, 18). It is difficult to understand how the AP2 complex works without clathrins in mediating endocytosis since, so far, endocytosis requiring AP2 but not clathrin has been reported only in *Aspergillus* (20, 23, 57). A significant finding in this study is that clathrins also function in Hb uptake. First, *Pf*CHC, *Pf*CLC, and *Pf*AP1 γ were localized at the collar of the cytostome. Second, *Pf*CHC KD resulted in abnormal long cytostome tubes, indicating *Pf*CHC is involved in the pinch-off of the cytostome, leading to reduced Hb digestion. Third, affinity pulldowns of clathrins also identified lower amounts of the *Pf*K13 complex (KIC3 or Eps15) and AP2 components, suggesting that the association may be loose or highly dynamic. Notably, studies in *T. gondii* have provided evidence supporting the localizations of clathrins and AP1 at the micropore (58–60). An EM report revealed a clathrin-like coat at the micropore surface (58). Two IFA studies revealed that both *Tg*CHC and *Tg*AP1 μ1 were localized at the PPM and *Tg*CHC was in the openings of IMC, the locations of micropores (59, 60) (See **Fig. S9** for details). All these indicate that clathrins are involved in the micropore process, although the exact relationships among K13, clathrins, AP2, and AP1 in *P. falciparum* and *T. gondii* are unknown and worthy of investigation in the future.

After cytostomes pinch off, Hb-containing vesicles, equivalent to early endosomes, either mature to become a food vacuole (FV) in the parasites at the early asexual developmental stage or fuse with existing FV in the parasites at the late stage (56, 61, 62). Several factors, including actin-myosin, dynamin, SNARE, Rab5a, and VPS45, were found to be involved in these processes (56, 62–64). Inhibition of actin destabilization and the activities of dynamin, myosin, and SNARE resulted in Hb-filled long cytostome tubes, whereas inhibition of actin stabilization and mislocalization of VPS45 caused an accumulation of Hb-containing vesicles in the parasite, indicating that these factors are needed for the pinch-off of Hb-containing vesicles and their subsequent transport to the FV, respectively (56, 62–64). The similar Hb-filled long cytostome tubes after *Pf*CHC KD suggest that clathrins are also involved in cytostome pinch-off in *P. falciparum*.

Our EM analysis showed CCV-like polygonal cells in the CCVs purified from parasites. However, the thickness of the electron-dense coat of CCVs is significantly thinner than CCVs in mammalian cells (7, 42), probably due to the highly diverse domain structures of clathrin and AP subunits in the parasite. *Pf*CHC KD caused a significant defect in merozoite invasion, indicating that clathrin is critical for this process. Previous studies showed that *Pf*AP1 µ1 was colocalized with rhoptry (16) and clathrin-based transport was found to be vital for exporting proteins from Golgi to invasion-related apical secretory organelle (rhoptry and microneme) in *T. gondii* (14, 59, 60). Thus, it is of great interest to elucidate the functions of clathrins in this secretory pathway.

## MATERIALS AND METHODS

### Parasite culture

*P. falciparum* clone 3D7 was cultured in type O^+^ red blood cells as described (65). Parasites were synchronized by treating the ring-stage parasites with 5% D-sorbitol.

### Phylogenetic comparison

GenBank entries of CHC, CLC, and AP1 subunits in model eukaryotes were retrieved for phylogenetic analysis. Sequence alignment and phylogenetic analysis were performed using the CLUSTAL Omega program (66).

### Genetic manipulation of *Pf*CHC, *Pf*CLC, and *Pf*AP1-**γ**

To tag the C-terminus of *Pf*CHC with GFP, a *PfCHC* fragment was amplified using primer pair 1 from the *P. falciparum* genomic DNA and cloned into a modified pBluescript SK plasmid to fuse with the GFP and pDT 3’ UTR (67, 68) (**Table S5**). This cassette was then subcloned into pHD22Y at the *Bam*HI and *Not*I sites to produce pHD22Y/*Pf*CHC-GFP (69). For APEX2 tagging, the *APEX2* fragment was amplified using primer pair 2 (**Table S5**). The GFP fragment in the pHD22Y/*Pf*CHC-GFP plasmid was replaced by the APEX2 tag to generate the pHD22Y/*Pf*CHC-APEX2. To tag the C-terminus of *Pf*CLC and *Pf*AP1 γ with the GFP or APEX2 tag, the *PfCLC,* and *Pf*AP1 γ fragments were amplified using primer pair 3 and 4, respectively (**Table S5**). pHD22Y/*Pf*CLC-GFP, pHD22Y/*Pf*CLC-APEX2, pHD22Y/*Pf*AP1 γ- GFP, and pHD22Y/AP1 γ-APEX2 were generated in the same ways as the *Pf*CHC GFP and APEX2 tagging. To knock down *PfCHC* using the TetR-DOZI system, the *Pf*CHC C-terminal fragment fused with GFP was amplified by using primer pair 5 using the pHD22Y/*Pf*CHC-GFP plasmid as a template and cloned into pMG75 upstream of the 10 X aptamer insertion at *BssH*II and *Dra*III sites to obtain the final construct pMG75/*Pf*CHC-GFP (**Table S5**). Parasite transfection was performed using an RBC loading method (70). Resistant parasite clones were obtained by limiting dilution (71). The correct integration was screened by integration-specific PCR using a forward primer upstream of the homologous recombination region and a reverse primer located in the GFP or APEX2 tag (**Table S5**).

### IPs and mass spectrometry

For GFP-tagged proteins, IPs were performed using the GFP-Trap agarose according to the manufacturer’s protocol (ChromoTek). The wild-type 3D7 parasites were used as IP controls. The elutes were separated briefly in an SDS-PAGE gel and proteins in the gel were excised and digested as described (72). The digests were analyzed by LC/MS/MS by the established methods using a Waters NanoAcquity HPLC system interfaced with a Q Exactive™ Hybrid Quadrupole- Orbitrap Mass Spectrometer (Thermo Scientific) (73–76). The mass spectrometry proteomics data have been deposited to the ProteomeXchange Consortium via the PRIDE (77) partner repository with the dataset identifier PXD031805 and 10.6019/PXD031805 (reviewer access login name: and password: Nd8UTnk1).

### Immunoblotting

To analyze the expression of *Pf*CHC, *Pf*CLC, and *Pf*AP1 γ in *Pf*CHC::GFP, *Pf*CLC::GFP, *Pf*AP1 γ::GFP, *Pf*CHC::APEX2, *Pf*CLC::APEX2, and TetR-CHC::GFP parasite lines, protein extracts from the respective parasite lines at different stages were analyzed by SDS-PAGE and immunoblotting using anti-GFP mAb (1:1000, Roche, 11814460001), anti-APEX2 antibodies (1:1000, Innovagen, PA-APEX2-100), respectively, as primary antibodies and horseradish peroxidase-conjugated IgG (1:3000) as the secondary antibodies. The results were visualized with the ECL detection system using X-ray film (Invitrogen, Cat# WP20005).

### Immunofluorescence assay (IFA)

Thin smears of iRBCs were prepared on glass slides and fixed with acetone for 10 seconds. Then, the slides were incubated with various antibodies separately, including ERD2 (rabbit, 1: 500), PfEMP1 (rabbit anti-ATS, 1 μg/ml), REX1 (rabbit, 1:5000), REX3 (rabbit, 1:1000, SBP1 (mouse, 1:1000), KAHRP (mouse mAb 18.2, 1:500), and FIKK4.2 (mouse, 1:500) (78) as primary antibodies and then incubated with FITC conjugated anti-rabbit or mouse IgG (1:1000) (Sigma, AP132F, AP124F) antibodies as secondary antibodies. The anti-PfEMP1 antiserum was raised against the peptide DITSSESEYEELDINDIC in the conserved acidic terminal sequence (ATS) of the 3D7 PfEMP1 (Proteintech Group), which has proved to detect all PfEMP1 (79).

For the colocalization study, the *Pf*CHC::GFP iRBCs were used with anti-GFP (mouse mAb 1:1000, Roche, 11814460001, rabbit ab6556, 1:1000), ERD2 (rabbit, 1: 500, MR4), and Exp2 (mouse mAb 7.7, 1:500) (80). The slides were observed under a Nikon Eclipse E600 microscope at 400 X magnification, and the pictures were taken with a Nikon digital camera DXM 1200 (Nikon Instruments Inc.).

### Super-resolution microscopy

For super-resolution fluorescence microscopy, *Pf*CHC::GFP iRBCs were stained with Bodipy-TR-C5-ceramide, mounted in the Antifade Mountant with DAPI (Molecular Probes, Eugene, OR, USA), and visualized using a Nikon N-SIM/N-STORM microscope. Three-dimensional (3D) reconstructions and isosurface models in volume images of reconstructed stacks were performed using the IMARIS version 7.7.2 software suite (Bitplane AG). For the clarity of display, deconvolved z stacks were reconstructed in 3D with interpolation.

### TEM, APEX-based EM

Conventional transmission EM was performed as described (67). The APEX2-based EM was conducted as described (39). The DAB reaction was induced by applying 0.5 mM H2O2 to the iRBC samples for 1 min at 4°C, followed by 2% OsO4 staining for 30 min on ice with or without uranyl acetate staining. Samples were imaged using a JEOL Electron Microscopy.

### Purification of CCVs

CCVs were purified using GFP-Trap magnetic beads (Chromoteck) from *Pf*CHC::GFP parasites or by ultracentrifugation coupled with sucrose/Ficoll gradient from wild-type 3D7 parasites (43, 81). Briefly, trophozoites were homogenized in MES buffer [100 mM MES, 1 mM EGTA, and 0.5 mM MgCl2 (pH 6.7)] and then centrifuged at 17,000 *g* for 25 min, and the supernatant was transferred to either incubate with GFP-Trap magnetic beads for 2 h or further centrifuged at 56,000 *g* for 1 h. The GFP beads were washed with MES buffer and ready for TEM analysis. The pellet from the above centrifugation (56,000 *g* for 1 h) was suspended in MES buffer with 12.5% (w/v) of sucrose and Ficoll, then spun at 43,000*g* for 1 h. The supernatant was transferred to new tubes and centrifuged at 100,000*g* for 2.5 h. The small part of the pellets was resuspended in MES buffer for observation by TEM after negative staining or cryofixation. The rest of the pellets (90%) were resuspended in lysis buffer (10 mM Tris-Cl pH 7.5, 150 mM NaCl, 0.5 mM EDTA, 0.5 % Nonidet P40 Substitute) for proteomic analysis.

### Negative staining and cryoEM analysis of purified CCVs

Negative staining was performed according to a standard protocol with uranyl acetate. Briefly, 3 μl of the sample was applied onto a glow-discharged carbon-coated grid (Ted Pella) and incubated for 1 min. The sample was blotted with filter paper and washed with 10 μl of 2% uranyl acetate three times. Thirty seconds after the final wash, the grid was blotted and air-dried for imaging. CryoEM grid was prepared by plunge-freezing with an FEI Mark IV Vitrobot (Thermo Fisher). Three μl of the sample were applied onto a glow-discharged thin continuous carbon film-coated lacey grid. After incubation in the chamber of Vitrobot at 8°C and 100% humidity for 1 min, the grid was blotted for 15 s and plunge-frozen in liquid ethane. Images for both negative staining and cryogenic grids were obtained with a 200 kV FEI TF20 electron microscopy. The images were recorded with TIETZ F415MP CCD camera at a magnification of 50,000 ×, corresponding to a pixel size of 4.41 Å on the specimen level. Particles were picked with gautomatch (https://www2.mrc-lmb.cam.ac.uk/research/locally-developed-software/zhang-software/), and 2D classifications were performed in RELION(82).

### Hemozoin content assays

The amount of hemozoin in iRBCs was estimated as previously described (83). Briefly, 1 × 10^9^ highly synchronized iRBCs at late trophozoite stages were lysed by ice-cold distilled water.

After centrifugation at 4000 rpm at 4°C for 30 min, the pellet was washed with ice-cold distilled water, dissolved in 1 ml of 0.1M NaOH, and incubated at 50°C for 10 min. The concentration of hemozoin in the supernatant was determined by measuring the absorbance at 400 nm.

### Hb content in parasites

Hb inside parasites was analyzed using an established method (84). First, the resealed RBCs were prepared in the presence of 50 μM fluorescein-dextran (anionic, 10 kDa; life-tech). Mature schizonts were added to the resealed RBCs. The green fluorescence signals in the parasite at the trophozoite stage (30 hpi) were examined by flow cytometric analysis after parasites were released from iRBCs by saponin lysis.

### Statistical analysis

Data is represented as mean ± standard deviation (SD) from three independent experiments. T-test was used for statistical comparison between growth phenotypes before and after *Pf*CHC KD. Unless indicated otherwise, *P*< 0.05 was taken as statistically significant.

### Data availability

All data was made publicly available on common data repositories. The mass spectrometry proteomics data have been deposited to the ProteomeXchange Consortium via the PRIDE partner repository with the dataset identifier PXD031805 and 10.6019/PXD031805.

## Acknowledgments

Anti-REX1, REX3, SBP1c, and aldolase antibodies were kind gifts from Dr. Tobias Spielmann. Anti-KAHRP, Exp2, and FIKK4.2 antibodies were obtained from the European Malaria Reagent Repository (EMRR: www.malariaresearch.eu) and were donated by Dr. Jana McBride and Dr. Odile Mercereau-Puijalon, respectively. Anti-ERD2 was obtained from The Malaria Research and Reference Reagent Resource Center (MR4). We want to thank Richard Jones at MS Bioworks and Dale Chaput at the USF Proteomics Core facility for their assistance with the proteomic analysis.

## References

1. Faini M, Beck R, Wieland FT, Briggs JA. Vesicle coats: structure, function, and general principles of assembly. Trends Cell Biol. 2013;23(6):279–88.

2. Kirchhausen T, Owen D, Harrison SC. Molecular structure, function, and dynamics of clathrin-mediated membrane traffic. Cold Spring Harb Perspect Biol. 2014;6(5):a016725.

3. Mettlen M, Chen PH, Srinivasan S, Danuser G, Schmid SL. Regulation of Clathrin-Mediated Endocytosis. Annu Rev Biochem. 2018;87:871–96.

4. Duncan MC. New directions for the clathrin adaptor AP-1 in cell biology and human disease. Current Opinion in Cell Biology. 2022;76.

5. Kirchhausen T. Three ways to make a vesicle. Nat Rev Mol Cell Biol. 2000;1(3):187–98.

6. McMahon HT, Mills IG. COP and clathrin-coated vesicle budding: different pathways, common approaches. Curr Opin Cell Biol. 2004;16(4):379–91.

7. Robinson MS. Forty years of clathrin-coated vesicles. Traffic. 2015;16(12):1210–38.

8. Bonifacino JS. Adaptor proteins involved in polarized sorting. J Cell Biol. 2014;204(1):7–17.

9. Nakatsu F, Hase K, Ohno H. The Role of the Clathrin Adaptor AP-1: Polarized Sorting and Beyond. Membranes (Basel). 2014;4(4):747–63.

10. Castillon GA, Burriat-Couleru P, Abegg D, Criado Santos N, Watanabe R. Clathrin and AP1 are required for apical sorting of glycosyl phosphatidyl inositol-anchored proteins in biosynthetic and recycling routes in Madin-Darby canine kidney cells. Traffic. 2018;19(3):215–28.

11. Gravotta D, Perez Bay A, Jonker CTH, Zager PJ, Benedicto I, Schreiner R, et al. Clathrin and clathrin adaptor AP-1 control apical trafficking of megalin in the biosynthetic and recycling routes. Mol Biol Cell. 2019;30(14):1716–28.

12. Edeling MA, Smith C, Owen D. Life of a clathrin coat: insights from clathrin and AP structures. Nat Rev Mol Cell Biol. 2006;7(1):32–44.

13. WHO. Would Malaria Report 2023. Geneva: WHO. 2023.

14. Tomavo S, Slomianny C, Meissner M, Carruthers VB. Protein trafficking through the endosomal system prepares intracellular parasites for a home invasion. PLoS Pathog. 2013;9(10):e1003629.

15. Klinger CM, Ramirez-Macias I, Herman EK, Turkewitz AP, Field MC, Dacks JB. Resolving the homology-function relationship through comparative genomics of membrane-trafficking machinery and parasite cell biology. Mol Biochem Parasitol. 2016;209(1-2):88–103.

16. Kaderi Kibria KM, Rawat K, Klinger CM, Datta G, Panchal M, Singh S, et al. A role for adaptor protein complex 1 in protein targeting to rhoptry organelles in Plasmodium falciparum. Biochim Biophys Acta. 2015;1853(3):699–710.

17. Henrici RC, Edwards RL, Zoltner M, van Schalkwyk DA, Hart MN, Mohring F, et al. The Plasmodium falciparum Artemisinin Susceptibility-Associated AP-2 Adaptin mu Subunit is Clathrin Independent and Essential for Schizont Maturation. mBio. 2020;11(1).

18. Birnbaum J, Scharf S, Schmidt S, Jonscher E, Hoeijmakers WAM, Flemming S, et al. A Kelch13-defined endocytosis pathway mediates artemisinin resistance in malaria parasites. Science. 2020;367(6473):51-9.

19. Yang J, Long S, Hide G, Lun ZR, Lai DH. Apicomplexa micropore: history, function, and formation. Trends Parasitol. 2024;40(5):416–26.

20. Spielmann T, Gras S, Sabitzki R, Meissner M. Endocytosis in Plasmodium and Toxoplasma Parasites. Trends Parasitol. 2020;36(6):520–32.

21. Xie SC, Ralph SA, Tilley L. K13, the Cytostome, and Artemisinin Resistance. Trends Parasitol. 2020;36(6):533–44.

22. Yang T, Yeoh LM, Tutor MV, Dixon MW, McMillan PJ, Xie SC, et al. Decreased K13 Abundance Reduces Hemoglobin Catabolism and Proteotoxic Stress, Underpinning Artemisinin Resistance. Cell Rep. 2019;29(9):2917–28 e5.

23. Behrens HM, Schmidt S, Spielmann T. The newly discovered role of endocytosis in artemisinin resistance. Med Res Rev. 2021;41(6):2998–3022.

24. Tutor Madel V. SGJ, Siddiqui Ghizal, Creek Darren J., Tilley Leann, Ralph Stuart A. The Plasmodium falciparum artemisinin resistance-associated protein Kelch 13 is required for formation of normal cytostomes 10.7554/eLife.90290.1. eLife 2023;12(5):305A.

25. Liffner B, Cepeda Diaz AK, Blauwkamp J, Anaguano D, Frolich S, Muralidharan V, et al. Atlas of Plasmodium falciparum intraerythrocytic development using expansion microscopy. Elife. 2023;12.

26. Wan W, Dong H, Lai DH, Yang J, He K, Tang X, et al. The Toxoplasma micropore mediates endocytosis for selective nutrient salvage from host cell compartments. Nat Commun. 2023;14(1):977.

27. Koreny L, Mercado-Saavedra BN, Klinger CM, Barylyuk K, Butterworth S, Hirst J, et al. Stable endocytic structures navigate the complex pellicle of apicomplexan parasites. Nat Commun. 2023;14(1):2167.

28. Goodman OB, Krupnick JG, Gurevich VV, Benovic JL, Keen JH. Arrestin/clathrin interaction - Localization of the arrestin binding locus to the clathrin terminal domain. Journal of Biological Chemistry. 1997;272(23):15017–22.

29. ter Haar E, Musacchio A, Harrison SC, Kirchhausen T. Atomic structure of clathrin: a beta propeller terminal domain joins an alpha zigzag linker. Cell. 1998;95(4):563–73.

30. ter Haar E, Harrison SC, Kirchhausen T. Peptide-in-groove interactions link target proteins to the beta-propeller of clathrin. Proc Natl Acad Sci U S A. 2000;97(3):1096–100.

31. Pishvaee B, Munn A, Payne GS. A novel structural model for regulation of clathrin function. Embo Journal. 1997;16(9):2227–39.

32. Brodsky FM. Diversity of Clathrin Function: New Tricks for an Old Protein. Annu Rev Cell Dev Bi. 2012;28:309–36.

33. Morris KL, Jones JR, Halebian M, Wu SP, Baker M, Armache JP, et al. Cryo-EM of multiple cage architectures reveals a universal mode of clathrin self-assembly. Nature Structural & Molecular Biology. 2019;26(10):890-+.

34. Teo G, Liu G, Zhang J, Nesvizhskii AI, Gingras AC, Choi H. SAINTexpress: improvements and additional features in Significance Analysis of INTeractome software. J Proteomics. 2014;100:37–43.

35. Hallee S, Boddey JA, Cowman AF, Richard D. Evidence that the Plasmodium falciparum Protein Sortilin Potentially Acts as an Escorter for the Trafficking of the Rhoptry-Associated Membrane Antigen to the Rhoptries. mSphere. 2018;3(1).

36. Hallée S, Counihan NA, Matthews K, de Koning-Ward TF, Richard D. The malaria parasite Sortilin is essential for merozoite formation and apical complex biogenesis. Cellular Microbiology. 2018;20(8).

37. Shadija N, Dass S, Xu W, Wang L, Ke H. Functionality of the V-type ATPase during asexual growth and development of Plasmodium falciparum. J Biol Chem. 2024;300(9):107608.

38. Alder A, Sanchez CP, Russell MRG, Collinson LM, Lanzer M, Blackman MJ, et al. The role of Plasmodium V-ATPase in vacuolar physiology and antimalarial drug uptake. Proc Natl Acad Sci U S A. 2023;120(30):e2306420120.

39. Martell JD, Deerinck TJ, Lam SS, Ellisman MH, Ting AY. Electron microscopy using the genetically encoded APEX2 tag in cultured mammalian cells. Nat Protoc. 2017;12(9):1792–816.

40. Zumthor JP, Cernikova L, Rout S, Kaech A, Faso C, Hehl AB. Static Clathrin Assemblies at the Peripheral Vacuole-Plasma Membrane Interface of the Parasitic Protozoan. Plos Pathogens. 2016;12(7).

41. Bannister LH, Hopkins JM, Fowler RE, Krishna S, Mitchell GH. Ultrastructure of rhoptry development in Plasmodium falciparum erythrocytic schizonts. Parasitology. 2000;121 ( Pt 3):273–87.

42. Tao-Cheng JH. Stimulation-induced differential redistributions of clathrin and clathrin-coated vesicles in axons compared to soma/dendrites. Mol Brain. 2020;13(1):141.

43. Paraan M, Mendez J, Sharum S, Kurtin D, He H, Stagg SM. The structures of natively assembled clathrin-coated vesicles. Sci Adv. 2020;6(30):eaba8397.

44. Zhang M, Wang C, Otto TD, Oberstaller J, Liao X, Adapa SR, et al. Uncovering the essential genes of the human malaria parasite Plasmodium falciparum by saturation mutagenesis. Science. 2018;360(6388).

45. Ganesan SM, Falla A, Goldfless SJ, Nasamu AS, Niles JC. Synthetic RNA-protein modules integrated with native translation mechanisms to control gene expression in malaria parasites. Nat Commun. 2016;7:10727.

46. Goldfless SJ, Wagner JC, Niles JC. Versatile control of Plasmodium falciparum gene expression with an inducible protein-RNA interaction. Nat Commun. 2014;5:5329.

47. Spillman NJ, Beck JR, Goldberg DE. Protein export into malaria parasite-infected erythrocytes: mechanisms and functional consequences. Annu Rev Biochem. 2015;84:813–41.

48. Jonsdottir TK, Gabriela M, Crabb BS, T FdK-W, Gilson PR. Defining the Essential Exportome of the Malaria Parasite. Trends Parasitol. 2021;37(7):664–75.

49. Krai P, Dalal S, Klemba M. Evidence for a Golgi-to-endosome protein sorting pathway in Plasmodium falciparum. PLoS One. 2014;9(2):e89771.

50. Quevillon E, Spielmann T, Brahimi K, Chattopadhyay D, Yeramian E, Langsley G. The Plasmodium falciparum family of Rab GTPases. Gene. 2003;306:13–25.

51. Kats LM, Cooke BM, Coppel RL, Black CG. Protein trafficking to apical organelles of malaria parasites - building an invasion machine. Traffic. 2008;9(2):176–86.

52. Van Wye J, Ghori N, Webster P, Mitschler RR, Elmendorf HG, Haldar K. Identification and localization of rab6, separation of rab6 from ERD2 and implications for an ’unstacked’ Golgi, in Plasmodium falciparum. Mol Biochem Parasitol. 1996;83(1):107–20.

53. Dasaradhi PVN, Korde R, Thompson JK, Tanwar C, Nag TC, Chauhan VS, et al. Food vacuole targeting and trafficking of falcipain-2, an important cysteine protease of human malaria parasite. Mol Biochem Parasit. 2007;156(1):12–23.

54. Subramanian S, Sijwali PS, Rosenthal PJ. Falcipain cysteine proteases require bipartite motifs for trafficking to the food vacuole. Journal of Biological Chemistry. 2007;282(34):24961–9.

55. Klemba M, Beatty W, Gluzman I, Goldberg DE. Trafficking of plasmepsin II to the food vacuole of the malaria parasite Plasmodium falciparum (vol 164, pg 47, 2004). Journal of Cell Biology. 2004;164(4):625-.

56. Abu Bakar N, Klonis N, Hanssen E, Chan C, Tilley L. Digestive-vacuole genesis and endocytic processes in the early intraerythrocytic stages of. Journal of Cell Science. 2010;123(3):441–50.

57. Martzoukou O, Amillis S, Zervakou A, Christoforidis S, Diallinas G. The AP-2 complex has a specialized clathrin-independent role in apical endocytosis and polar growth in fungi. Elife. 2017;6.

58. Nichols BA, Chiappino ML, Pavesio CEN. Endocytosis at the Micropore of Toxoplasma- Gondii. Parasitology Research. 1994;80(2):91–8.

59. Pieperhoff MS, Schmitt M, Ferguson DJ, Meissner M. The role of clathrin in post-Golgi trafficking in Toxoplasma gondii. PLoS One. 2013;8(10):e77620.

60. Venugopal K, Werkmeister E, Barois N, Saliou JM, Poncet A, Huot L, et al. Dual role of the Toxoplasma gondii clathrin adaptor AP1 in the sorting of rhoptry and microneme proteins and in parasite division. PLoS Pathog. 2017;13(4):e1006331.

61. Lazarus MD, Schneider TG, Taraschi TF. A new model for hemoglobin ingestion and transport by the human malaria parasite Plasmodium falciparum. J Cell Sci. 2008;121(11):1937–49.

62. Milani KJ, Schneider TG, Taraschi TF. Defining the morphology and mechanism of the hemoglobin transport pathway in Plasmodium falciparum-infected erythrocytes. Eukaryot Cell. 2015;14(4):415–26.

63. Elliott DA, McIntosh MT, Hosgood HD, 3rd, Chen S, Zhang G, Baevova P, et al. Four distinct pathways of hemoglobin uptake in the malaria parasite Plasmodium falciparum. Proc Natl Acad Sci U S A. 2008;105(7):2463–8.

64. Jonscher E, Flemming S, Schmitt M, Sabitzki R, Reichard N, Birnbaum J, et al. PfVPS45 Is Required for Host Cell Cytosol Uptake by Malaria Blood Stage Parasites. Cell Host & Microbe. 2019;25(1):166-+.

65. Trager W, Jensen JB. Human malaria parasites in continuous culture. Science. 1976;193(4254):673–5.

66. Madeira F, Madhusoodanan N, Lee JH, Eusebi A, Niewielska A, Tivey ARN, et al. The EMBL-EBI Job Dispatcher sequence analysis tools framework in 2024. Nucleic Acids Research. 2024;52(W1):W521–W5.

67. Miao J, Fan Q, Cui L, Li X, Wang H, Ning G, et al. The MYST family histone acetyltransferase regulates gene expression and cell cycle in malaria parasite Plasmodium falciparum. Mol Microbiol. 2010;78(4):883–902.

68. Fan Q, Miao J, Cui L, Cui L. Characterization of PRMT1 from Plasmodium falciparum. Biochem J. 2009;421(1):107–18.

69. Fidock DA, Wellems TE. Transformation with human dihydrofolate reductase renders malaria parasites insensitive to WR99210 but does not affect the intrinsic activity of proguanil. Proc Natl Acad Sci U S A. 1997;94(20):10931–6.

70. Deitsch K, Driskill C, Wellems T. Transformation of malaria parasites by the spontaneous uptake and expression of DNA from human erythrocytes. Nucleic Acids Res. 2001;29(3):850–3.

71. Rosario V. Cloning of naturally occurring mixed infections of malaria parasites. Science. 1981;212(4498):1037–8.

72. Lasonder E, Ishihama Y, Andersen JS, Vermunt AM, Pain A, Sauerwein RW, et al. Analysis of the Plasmodium falciparum proteome by high-accuracy mass spectrometry. Nature. 2002;419(6906):537-42.

73. Miao J, Chen Z, Wang Z, Shrestha S, Li X, Li R, et al. Sex-Specific Biology of the Human Malaria Parasite Revealed from the Proteomes of Mature Male and Female Gametocytes. Mol Cell Proteomics. 2017;16(4):537–51.

74. Kalamuddin M, Shakri AR, Wang C, Min H, Li X, Cui L, et al. MYST regulates DNA repair and forms a NuA4-like complex in the malaria parasite Plasmodium falciparum. mSphere. 2024:e0014024.

75. Miao J, Wang CQ, Lucky AB, Liang XY, Min H, Adapa SR, et al. A unique GCN5 histone acetyltransferase complex controls erythrocyte invasion and virulence in the malaria parasite Plasmodium falciparum. Plos Pathogens. 2021;17(8).

76. Lucky AB, Wang CQ, Liu M, Liang XY, Min H, Fan Q, et al. A type II protein arginine methyltransferase regulates merozoite invasion in Plasmodium falciparum. Commun Biol. 2023;6(1).

77. Perez-Riverol Y, Bai J, Bandla C, Garcia-Seisdedos D, Hewapathirana S, Kamatchinathan S, et al. The PRIDE database resources in 2022: a hub for mass spectrometry-based proteomics evidences. Nucleic Acids Res. 2022;50(D1):D543–D52.

78. Kats LM, Fernandez KM, Glenister FK, Herrmann S, Buckingham DW, Siddiqui G, et al. An exported kinase (FIKK4.2) that mediates virulence-associated changes in Plasmodium falciparum-infected red blood cells. Int J Parasitol. 2014;44(5):319–28.

79. Sharling L, Sowa KM, Thompson J, Kyriacou HM, Arnot DE. Rapid and specific biotin labelling of the erythrocyte surface antigens of both cultured and ex-vivo Plasmodium parasites. Malar J. 2007;6:66.

80. Hall R, McBride J, Morgan G, Tait A, Zolg JW, Walliker D, et al. Antigens of the erythrocytes stages of the human malaria parasite Plasmodium falciparum detected by monoclonal antibodies. Mol Biochem Parasitol. 1983;7(3):247–65.

81. Bouvier D, Tremblay ME, Riad M, Corera AT, Gingras D, Horn KE, et al. EphA4 is localized in clathrin-coated and synaptic vesicles in adult mouse brain. J Neurochem. 2010;113(1):153–65.

82. Scheres SHW. Processing of Structurally Heterogeneous Cryo-EM Data in RELION. Method Enzymol. 2016;579:125–57.

83. Waldecker M, Dasanna AK, Lansche C, Linke M, Srismith S, Cyrklaff M, et al. Differential time-dependent volumetric and surface area changes and delayed induction of new permeation pathways in P. falciparum-infected hemoglobinopathic erythrocytes. Cell Microbiol. 2017;19(2).

84. Klonis N, Crespo-Ortiz MP, Bottova I, Abu-Bakar N, Kenny S, Rosenthal PJ, et al. Artemisinin activity against Plasmodium falciparum requires hemoglobin uptake and digestion. Proc Natl Acad Sci U S A. 2011;108(28):11405–10.

